# MicroRNA-138 negatively regulates the hypoxia-inducible factor 1α to suppress melanoma growth and metastasis

**DOI:** 10.1101/566224

**Authors:** Haijiang Qiu, Fangchao Chen, Minjun Chen

## Abstract

Melanoma with rapid progression towards metastasis becomes the deadliest form of skin cancer. However, the mechanism of melanoma growth and metastasis is still unclear. Here, we found that miRNA-138 was low expression and hypoxia-inducible factor 1α (HIF1α) was high expression in the patient’s melanoma tissue, and they had a significant negative correlation (r = -0.937, *P* < 0.001). Patients with miRNA-138^low^/HIF1α^high^ signature were predominant in late stage. Further, bioinformatic analysis demonstrated that miRNA-138 directly targeted HIF1α. We found that the introduction of miRNA-138 mimics to A375 cells could reduced HIF1α mRNA expression, and suppressed the cell proliferation, migration and invasion. Overexpression of miRNA-138 or inhibition of HIF1α significantly suppressed the growth and metastasis of melanoma *in vivo*. Our study demonstrates the role and clinical relevance of miRNA-138 and HIF1α in melanoma cell growth and metastasis, providing a novel therapeutic target for suppression of melanoma growth and metastasis.

## INTRODUCTION

Melanomas, the most common and aggressive form of skin cancer, are highly heterogeneous and can progress rapidly to metastatic disease (Gupta et al., 2005; Luo et al., 2016). However, the mechanism and effective therapies for melanoma remain elusive. Recently, many studies have pointed toward MicroRNAs (miRNAs) as playing key roles in the development, metastasis and prognosis of melanoma (Deng et al., 2016; Hanniford et al., 2015). miRNAs, 18-25 nucleotides in length non-coding RNAs, act as negative regulators of post-transcriptional gene regulation through directly binding to the 3’-untranslational region (UTR) of their target mRNAs, and promoting translational suppression or their degradation, finally modulate the expression and function of those genes.

Studies have shown that miRNA-21 (Saldanha et al., 2016), miRNA-125 (Glud et al., 2010), miRNA-137 (Bemis et al., 2008), miRNA-142 (Jayawardana et al., 2016), miRNA-145 (Noguchi et al., 2012), miRNA-146b (Tembe et al., 2015), miRNA-155 (Levati et al., 2009), miRNA-205 (Dar et al., 2011), miRNA-182 (Segura et al., 2009), miRNA-193b (Chen et al., 2011), miRNA-342, and miRNA-608 (Jiao et al., 2018) are involved in the regulation of cell cycle and proliferation, invasion and migration, and cell survival or as diagnostic and prognostic biomarkers in melanoma. miRNA-138 is downregulated in a wide range of cancers including glioblastoma (Qiu et al., 2013), non-small cell lung cancer (Zhang et al., 2013), renal cell carcinoma (Yamasaki et al., 2012) and cholangiocarcinoma (Wang et al., 2013). miRNA-138 suppressed cancer cell invasion and migration by repressing H2AX (Wang et al., 2011), EZH2 (Qiu et al., 2013), Vimentin (Yamasaki et al., 2012), S100A1 (Sen et al., 2013), YAP1 (Xiao et al., 2016). However, the expression of miRNA-138 and its role in melanoma cell invasion and metastasis are still poorly understood.

In melanoma, HIF1α is well substantiated to promote tumorigenesis (Loftus et al., 2017). Elevated expression of HIF1α correlates with a cell phenotypic transition from proliferation to metastasis (O’Connell et al., 2013; Widmer et al., 2013). Additionally, mouse melanoma tumors exposed to hypoxia show increased tumor growth and metastatic potential (Cheli et al., 2012). These imply a crucial *in vivo* role for HIF1α in regulating the melanoma cell changes from proliferative to metastasis that lead to regional and distal tumor progression.

Here, we found that the expression of miRNA-138 was significantly down-regulated in both human melanoma tissues (HMTs) and melanoma cell lines. HIF1α has been identified as the direct downstream target of miRNA-138 in melanoma. Further studies showed that exogenous overexpression of miRNA-138 or knockdown HIF1α inhibits melanoma cell growth and metastasis, both *in vitro* and *in vivo*. Thus, miRNA-138/HIF1α is a potential target for the clinical diagnosis and treatment of melanoma.

## RESULTS

### miRNA-138 expression was downregulated in HMTs

Firstly, we quantitatively detected the expression of miRNA-138 and a variety of its related downstream proteins including FOXC1, SOX4, HIF1α, CDK6, E2F2, and E2F3 in HMTs and paracancerous tissues (PTs). The results showed that miRNA-138 was significantly downregulated in HMTs compared with PTs (Fig. 1A). Conversely, HIF1α had higher level than other downstream proteins in HMTs compared with PTs (Fig. 1B). Then, we analyzed the correlation between miRNA-138 and HIF1α using Pearson correlation coefficient. The results showed that the miRNA-138 level was negatively correlated with the HIF1α mRNA level (Fig. 1C). We further examined the difference of the miRNA-138 and HIF1α expression at different stages of melanoma. The late-stages of melanoma tissues had further decreased miRNA-138 and increased HIF1α expression (Fig. 1D). These results suggest that miRNA-138 and HIF1α may be involved in the growth and metastasis of melanoma and play opposite roles.

**Fig. 1.**
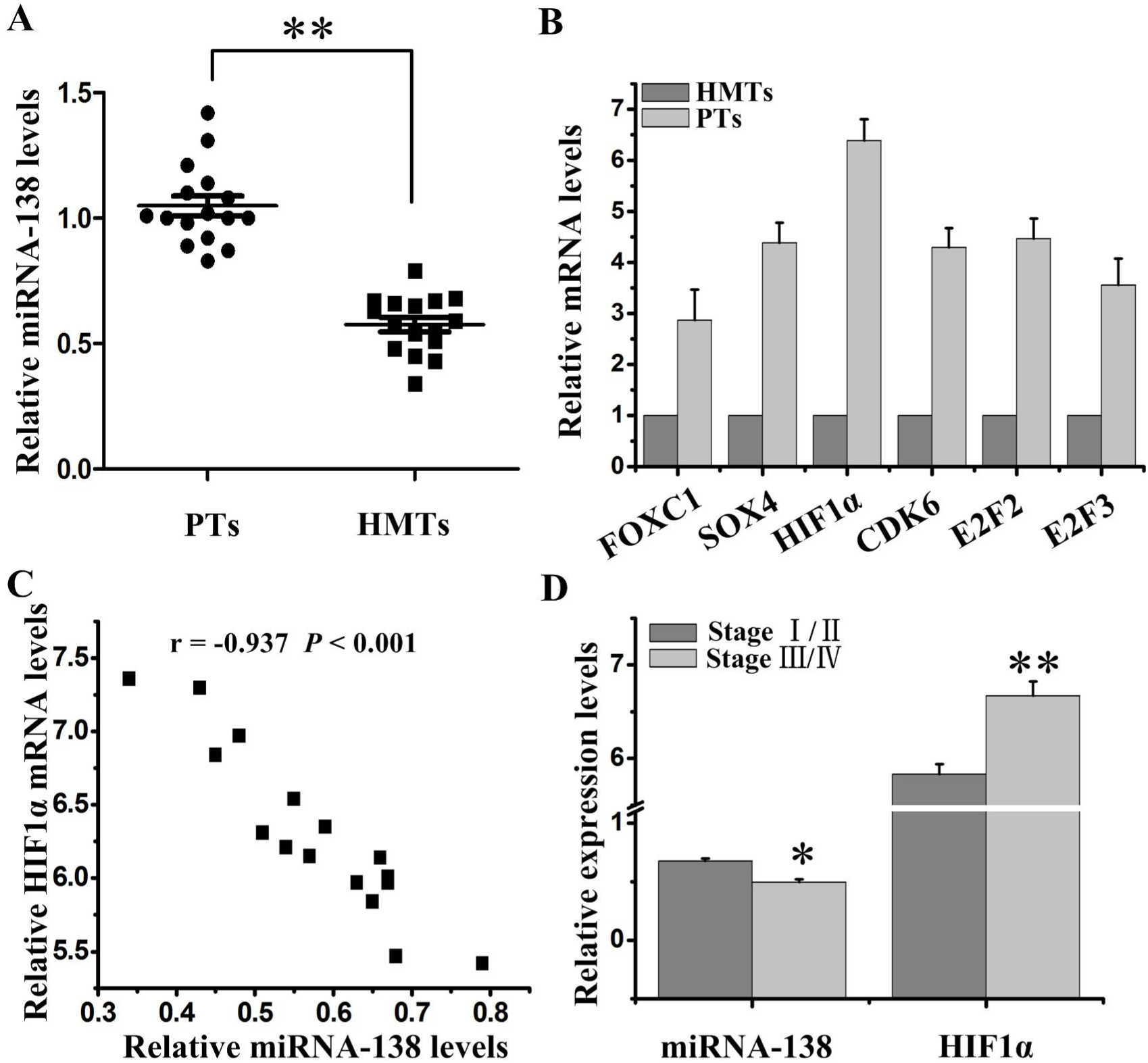
Expression of miRNA-138 and HIF1α in HMTs. (A) miRNA-138 expression levels in HMTs and PTs were assessed by qRT-PCR. (B) The mRNA expressions of FOXC1, SOX4, HIF1α, CDK6, E2F2, and E2F3 in HMTs and PTs were analyzed by qRT-PCR. (C) The correlation between miRNA-138 expression level and HIF1α mRNA level was estimated using Pearson’s correlation analysis. (D) The difference of miRNA-138 and HIF1α mRNA expression in stage I/II and III/IV of HMTs. \**P* < 0.05 and \*\**P* < 0.01 vs. control.

### miRNA-138 inhibited melanoma cell proliferation and invasion, and promoted cell apoptosis

The above results motivated us to observe whether miRNA-138 is involved in the growth and metastasis of melanoma. We designed loss-of-function and gain-of-function experiments to investigate the role of miRNA-138 in melanoma cells. We firstly observed that primary WM35 cells had low proliferation (Fig. 2A), migration (Fig. 2C), and invasion (Fig. 2D). However, proliferation, migration and invasion of miRNA-138 knockdown (KD) WM35 cells were increased. Metastatic A375 cells had high proliferation (Fig. 2B), migration (Fig. 2E), and invasion (Fig. 2F) However, the proliferation, migration and invasion of miRNA-138 overexpressing (OE) A375 cells were decreased. Those results suggested that miRNA-138 regulates the proliferation and invasion of melanoma cells.

**Fig. 2.**
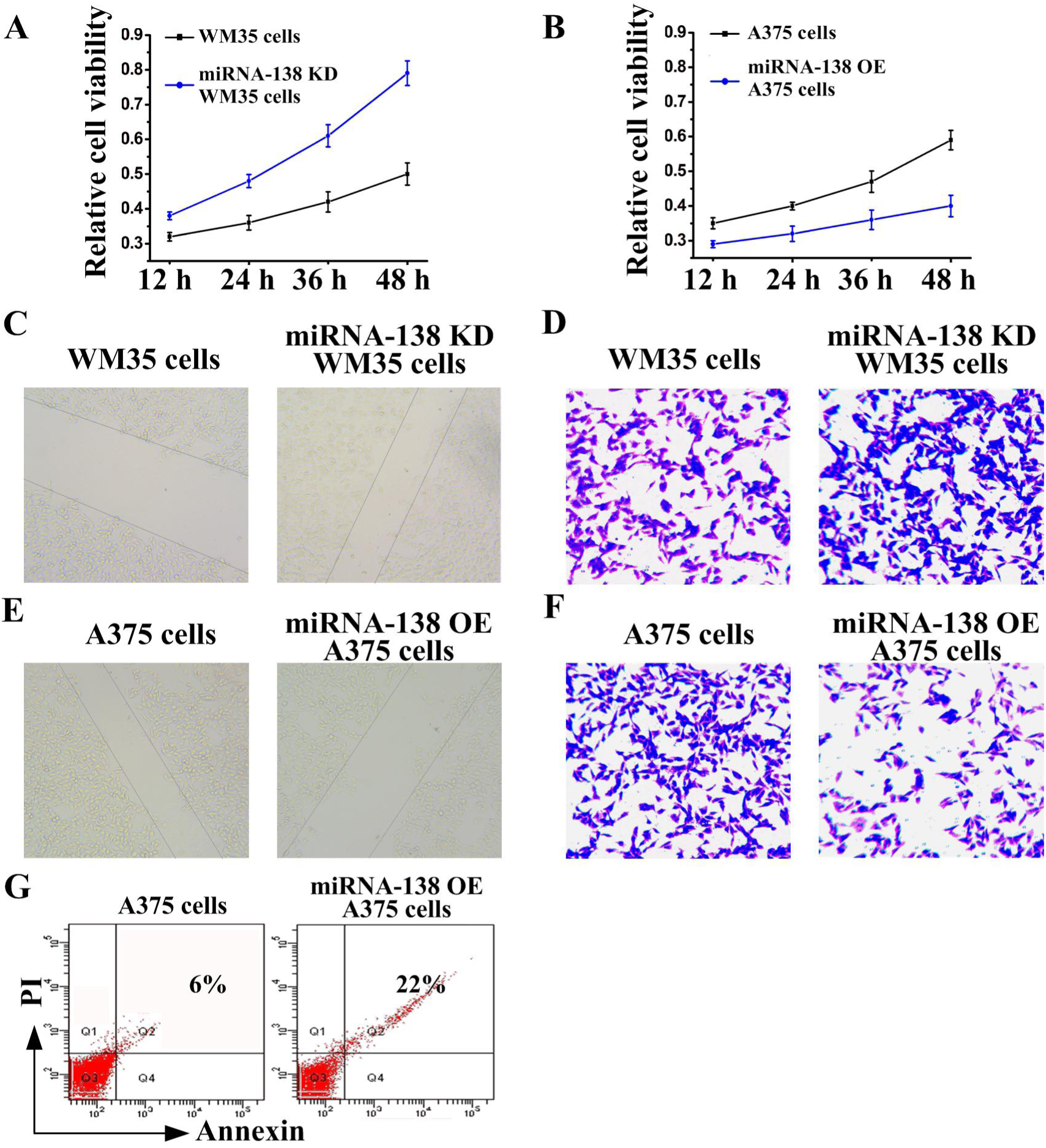
miRNA-138 regulates the growth and apoptosis of melanoma cells. (A, B) MTT assay, (C, E) wound healing assay and (D, F) transwell assay were employed to examine miRNA-138 knowdown (KD) WM35 cell and miRNA-138 overexpressing (OE) A375 cell proliferation, migration, and invasion, respectively. (G) Cell apoptosis rates of A375 cells following overexpression of miRNA-138 were detected by flow cytometry.

Then, we further observed the effect of miRNA-138 on apoptosis of melanoma cells. As shown in Figure 2G, cell apoptosis increased significantly in miRNA-138 overexpressing A375 cells compared with A375 cells. Therefore, miRNA-138 functioned as a suppressor of melanoma occurrence and development.

### HIF1α suppressed the effects of miRNA-138 on melanoma cells

Because HIF1α and miRNA-138 are negatively correlated, we wondered to explore whether HIF1α and miRNA-138 had an opposite effect. To this end, we applied siRNA knock down experiment. The results showed that siRNA HIF1α (si-HIF1α) significantly inhibited melanoma proliferation (Fig. 3A), migration (Fig. 3C), and invasion (Fig. 3D) in WM35 cells. Additionally, the effect of knockdown miRNA-138 on cell proliferation (Fig. 3B), migration (Fig. 3E), invasion (Fig. 3F), and apoptosis (Fig. 3G) was reversed by siRNA HIF1α in WM35 cells, suggesting that HIF1α and miRNA-138 are antagonistic in function.

**Fig. 3.**
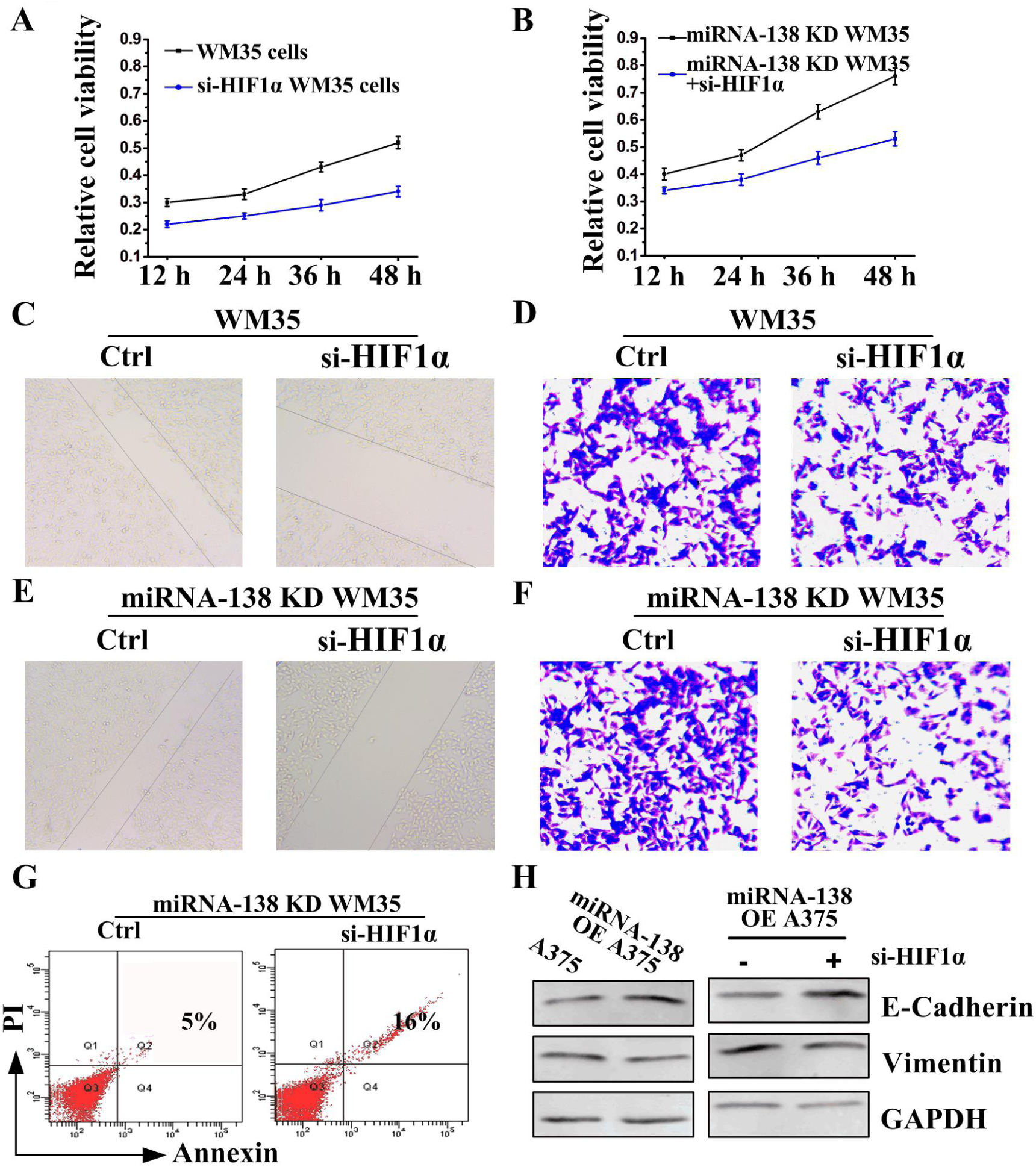
HIF1α also regulates the growth and apoptosis of melanoma cells. (A, B) MTT assay, (C, E) wound healing assay and (D, F) transwell assay were employed to examine WM35 or miRNA-138 knowdown (KD) WM35 cell proliferation, migration, and invasion, respectively, in presence of HIF1α siRNA (si-HIF1α). (G) Cell apoptosis rates of miRNA-138 knowdown WM35 cells following transfection si-HIF1α were detected by flow cytometry. (H) Western blot assay to detect E-cadherin and Vimentin expressions following si-HIF1α treatment after 48 h in A375 or miRNA-138 overexpressing A375 cells.

miRNA-138 played a regulatory role for epithelial to mesenchymal transition (EMT) (Zhang et al., 2016), which has an increased incidence of tumor metastasis (Qin et al., 2017). Thus, the effect of miRNA-138 on EMT was further examined. We found that miRNA-138 significantly promoted E-cadherin expression and inhibited Vimentin expression (Fig. 3H left), suggesting that the process of EMT was inhibited by miRNA-138. Importantly, siRNA HIF1α further promoted the effect of miRNA-138 (Fig. 3H right). These results indicate that functions of HIF1α and miRNA-138 were antagonistic in regulating proliferation, invasion and metastasis of melanoma cells.

### HIF1α mRNA was a direct target of miRNA-138

miRNA usually binds mRNA 3’UTR regions to suppress their translation (Deng et al., 2016). To further explore whether miRNA-138 have a direct binding with HIF1α mRNA and then downregulated the HIF1α expression, 3’UTR luciferase reporter plasmid assay was employed. The plasmid was constructed containing the mutant or wild-type miRNA-138-binding sequences of HIF1α 3’UTRs (Fig. 4A). miRNA-138 significantly inhibited the luciferase activities of WT HIF1α 3’UTR constructs (Fig. 4B), suggesting miRNA-138 directly targeted HIF1α.

**Fig. 4.**
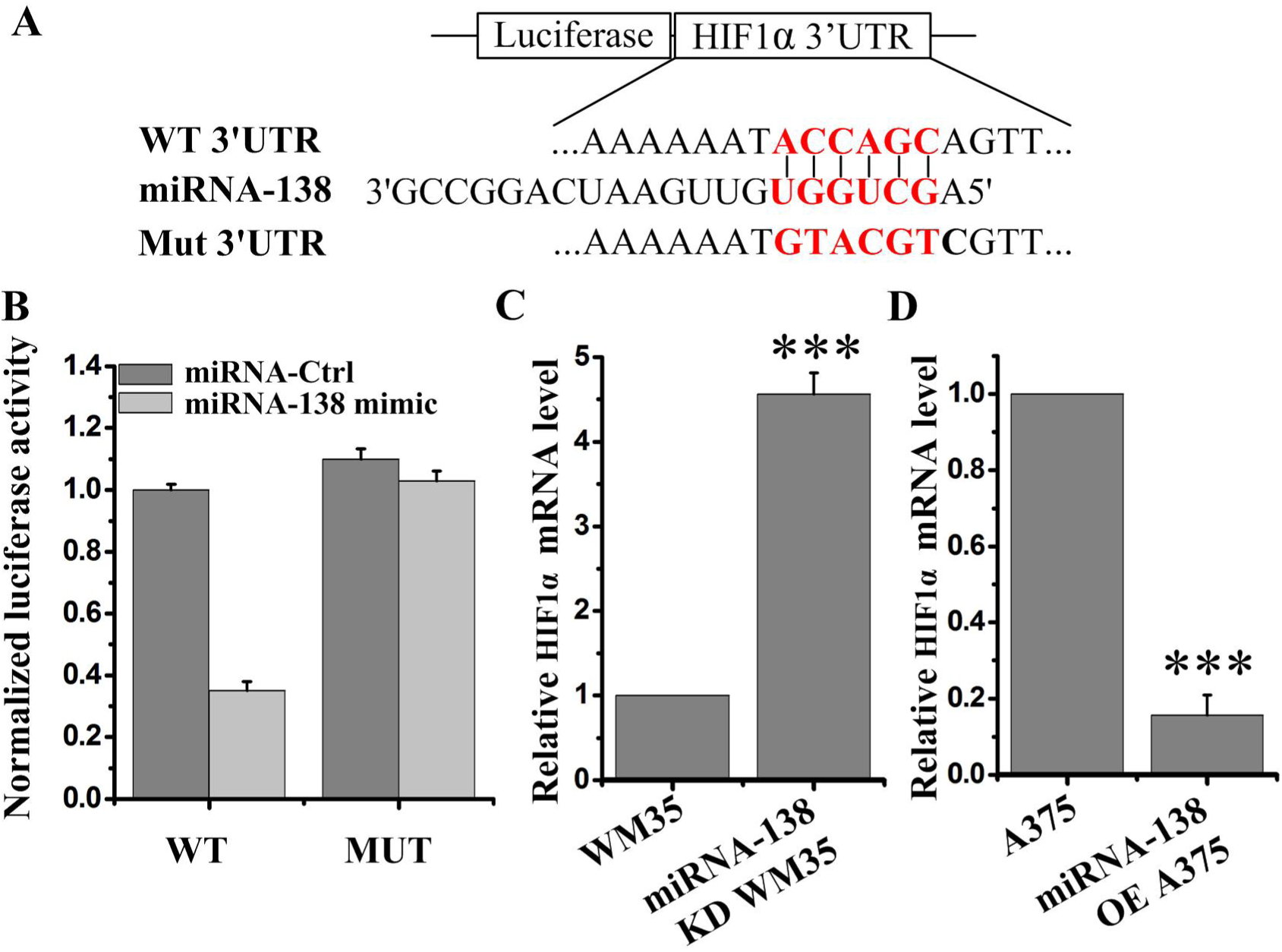
miRNA-138 directly targets HIF1α and degrades its mRNA. (A) The HIF1α 3′UTR region containing the wild type (WT) or mutant (MUT) binding site for miRNA-138. (B) The relative luciferase activity of HIF1α wild type or mutant 3’UTR in HEK293 cells following transfection with miRNA-138 mimic or miRNA-Control. (C-D) The HIF1α mRNA expression in miRNA-138 knowdown (KD) WM35 cells (C) and miRNA-138 overexpressing (OE) A375 cells (D) was detected by qRT-PCR. \*\*\**P* < 0.001 vs. control.

Subsequently, the changes of HIF1α mRNA levels under miRNA-138 knockdown or overexpression conditions were undertook to analyze by quantitative PCR. We found that miRNA-138 significantly reduced HIF1α mRNA levels (Fig.4C and 4D). These data indicate that miRNA-138 directly suppresses HIF1α translation to result in the decrease of its mRNA levels.

### miRNA-138 and HIF1α inhibitor synergistically suppressed melanoma growth and metastasis

To further observe the therapeutic effects of miRNA-138 and HIF1α *in vivo*, we used miRNA-138 knockdown and overexpression cells to inoculate Balb/c mice. The results showed that miRNA-138 knockdown WM35 tumors grew faster than primary WM35 tumors (Fig. 5A). Conversely, miRNA-138 overexpressing A375 tumors grew very slowly (Fig. 5B). Additionally, inhibition of HIF1α through oral administration of BAY 87-2243 also significantly inhibited the tumor growth (Fig. 5B) and metastasis (Fig. 5C and 5D). To our surprise, the combination therapy of overexpressing miRNA-138 and inhibition of HIF1α eradicated tumor growth and largely inhibited tumor metastasis, suggesting that the targeted therapy via miRNA-138 and HIF1α was a novel approach to melanoma treatment. The body weight of mice under the above treatment was no significant change (Fig. 5E), suggesting that the use of drugs in mice is reasonable.

**Fig. 5.**
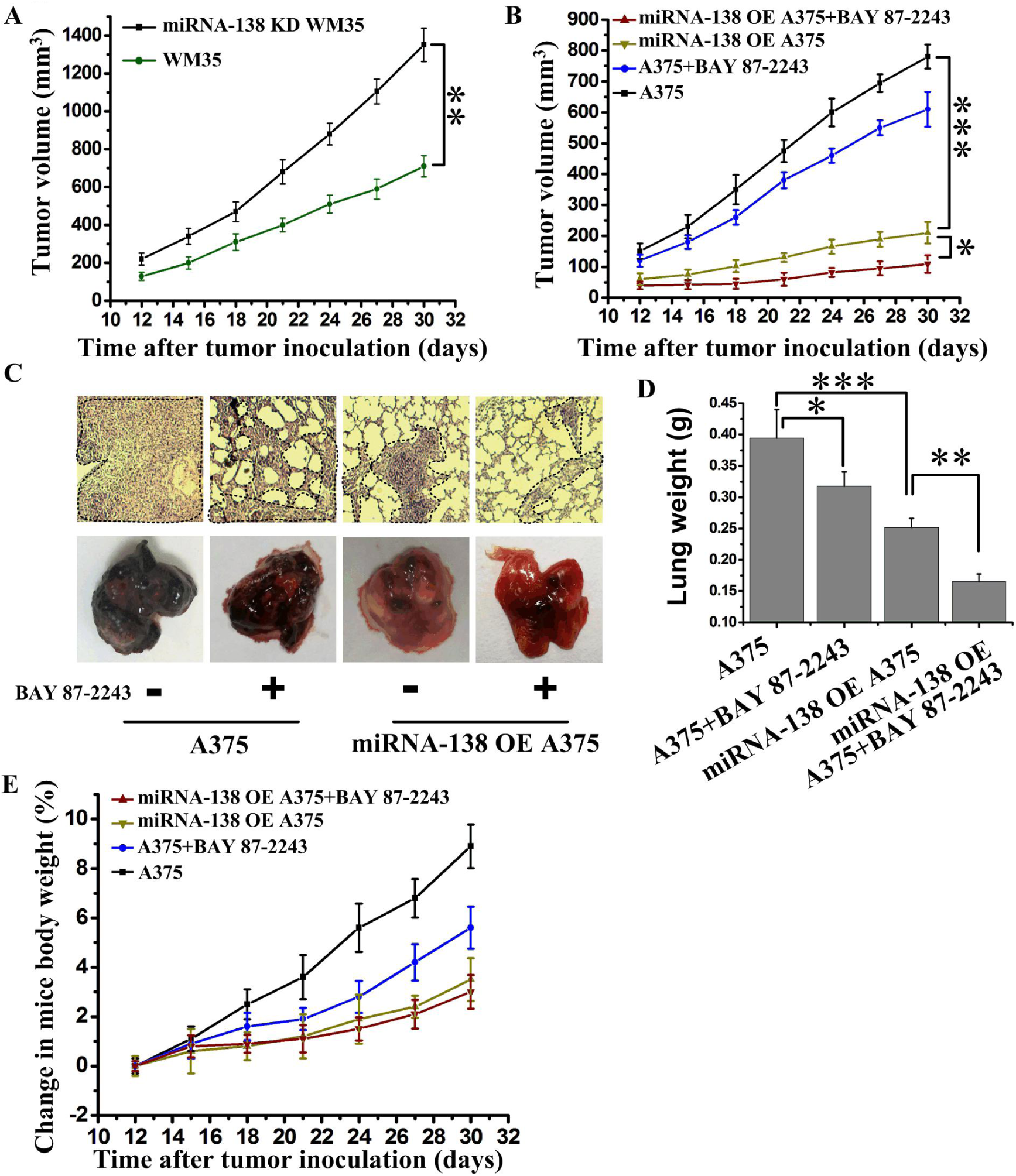
The upregulation of miRNA-138 and HIF1α inhibitor synergistically suppressed melanoma growth and metastasis. (A) Tumor growth curve from BALB/c mice with incubation of WM35 or miRNA-138 knowdown (KD) WM35 cells (n = 7 mice per group). (B) Tumor growth curve from BALB/c mice with incubation of A375 or miRNA-138 overexpressing (OE) A375 cells. BAY 87-2243 were orally administered as a 2-week continuous infusion at 9 mg/kg/day beginning on the fourth day after incubation (n = 6 mice per group). (C) Representative images of tumor metastases in BALB/c mice after i.v. injections of A375 or miRNA-138 overexpressing A375 cells on the twenty-fifth day. BAY 87-2243 were orally administered as a 2-week continuous infusion at 9 mg/kg/day beginning on the fourth day after incubation. Circles indicate lung metastases. (D) Quantitative analysis of the entire weight of lungs from (C). (E) Analysis of weight change in BALB/c mice from (B) (n = 6 mice per group). *P < 0.05, **P < 0.01, ***P < 0.001 vs. indicated group.

## DISCUSSION

miRNAs play more critical roles in the tumor development and prognosis (Deng et al., 2016). The function of miRNA-138 has been extensively studied in other tumors (Jiang et al., 2016; Yeh et al., 2013; Yu et al., 2015), but its role in melanoma tissue remains unclear. Herein, we identified and verified miRNA-138 as an effective tumor suppressor on cell migration, invasion and metastasis in melanoma. Our results also showed that the molecular mechanism underlying miRNA-138 action in melanoma inhibited cell proliferation and metastasis by directly targeting degradation of HIF1α. Tumor cells can adapt and survive under hypoxic conditions, this may be caused by HIF1-induced numerous target genes (Mineo et al., 2016), which mediate coagulation, angiogenesis, and metabolism to provide more nutrients and oxygen for tumor tissues (Ebersole et al., 2018). Studies have showed that HIF1α, a subunit of HIF1, can directly promote the expression of EMT-related genes (Jiang et al., 2011; Yang et al., 2008), enhancing tumor cell proliferation and metastasis. Indeed, HIF1α was high expression in melanoma tissues (Fig. 1B and 1D). Decreased HIF1α activity by RNA interference significantly ininhibited the E-cadherin expression and increased the vimentin expression, and reversed the process of EMT (Fig. 3H), suggesting that HIF1α may be a new target to treat melanoma metastasis. Other studies showed that HIF1α activated EMT regulators including SIP1, ZEB1 SLUG, SNAIL, or TWIST to promote of cancer cell metastasis (Huang et al., 2009; Lin et al., 2016; Yang et al., 2008; Yoo et al., 2011). In melanoma, the downstream signal regulation mechanism of HIF1α is still under our further study.

Current therapeutic interventions for highly metastatic melanoma are not satisfactory. Thus, efforts to understand the molecular mechanisms underlying the metastatic process of melanoma are a highly significant clinical importance. Our data from the tumor tissues of melanoma patients with different stages showed that the expression of miRNA-138 decreased progressively with the enhancement of metastatic potential and the corresponding HIF1α level increased gradually. Therefore, revealing the functions of miRNA-138 and HIF1α, and their correlation are important for the clinical treatment of melanoma. In our study, the combination therapy of overexpressing miRNA-138 and inhibition of HIF1α *in vivo* showed tumor growth and metastasis were eradicated (Fig.5B and 5C). Those results revealed the mechanism of melanoma invasion and metastasis.

To summarize, we identified miRNA-138 as a regulator for controlling melanoma cell proliferation, invasion and metastasis, and revealed a novel mechanism by which miRNA-138 negatively regulates the HIF1α to suppress melanoma growth and metastasis. Upregulation of the miRNA-138 may serve as a new therapeutic target for patients with melanoma.

## MATERIALS AND METHODS

### Tissue collection

Human HMTs (n = 16) were collected following surgical resection of cutaneous melanomas at the Second Affiliated Hospital of South China University of Technology from September 2017 to July 2018, following a protocol approved by the Hospital’s Ethics Committee. This cohort included 9 male and 7 female patients. The mean age was 59.6 years (range: 32-69 years). Tumor stages were determined according to the consensus of the European Thyroid Association (ETA) risk stratification system and the tumor, node, metastasis (TNM) staging system, respectively. Tissue specimens are diagnosed as cutaneous melanomas by histopathology and cytology.

### Cell lines and reagents

WM35 and A375 melanoma cells and HEK293 cells were purchased from the Cell Bank of Type Culture Collection of the Chinese Academy of Sciences (Shanghai, China) and cultured in Dulbecco’s Modified Eagle’s Medium (DMEM) supplemented with 10% fetal bovine serum (Sijiqing, hangzhou, China) at 37°C in a humidified atmosphere containing 5% CO_2_.

BAY 87-2243 (S7309) were purchased from Selleckchem. GAPDH (sc-32233) was obtained from Santa Cruz Biotechnology. The antibodies recognizing E-Cadherin (#14472) and Vimentin (#3390) were purchased from Cell Signaling Technology.

### Lentivirus production and transfection

miRNA-138 oligonucleotide mimics/inhibitors and their non-specific control were synthesized by GenePharma (Shanghai, China). Those sequences were inserted into the lentivirus vector pLL3.7 (Sigma) to generate a miRNA-138 mimic oligonucleotide vector (LV-miRNA-138) and its non-specific control vector (LV-NC), and a miRNA-138 inhibitor oligonucleotide vector (LV-miRNA-138 inhibitor) and its non-specific control vector (LV-NC inhibitor). The miRNA-138 overexpressing stable cells were established by infecting A375 cells with lentiviruses encoding LV-miRNA-138 and LV-NC followed by selection in 2 mg/ml puromycin. Similarly, the miRNA-138 knockdown stable cells were established by infecting WM35 cells with lentiviruses encoding LV-miRNA-138 inhibitor and LV-NC inhibitor followed by selection in 2 mg/ml puromycin. The overexpression and knockdown efficiency of miRNA-138 was detected by qPCR. HIF1α small interfering RNA (siRNA) (5’-CUAUGAACAUAAAGUCUGCTT-3’) or its non-specific control (5’-UUCUCCGAACGUGUCACGUTT-3’) were transfected into cells using Lipofectamine 3000 reagent (Invitrogen, Carlsbad, CA).

### Quantitative real-time PCR (qPCR) for genes and miRNAs

Total RNA and miRNA were isolated from HMTs using TRIzol (Invitrogen) according the manufacturer’s instructions and cDNA was generated using ReverTra Ace PCR RTMaster Mix with gDNA Remover (Toyobo). qPCR was performed using a TaqMan miRNA Assay according to manufacturer’s protocol (Applied Biosystems) by cDNA amplification: 40 cycles of 15 s at 95°C and 1 min at 60°C, and specific PCR products were generated using the following primers: 5’- AGCTGGTGTTGTGAATCAGGCCG-3’ (forward) and 5’-TGGTGTCGTGGAGTCG-3’ (reverse) were used for detecting miRNA-138; 5’- CTCGCTTCGGCAGCACA-3’ (forward) and 5’-AACGCTTCACGAATTTGCGT-3’ (reverse) were used for detecting U6; 5’-GCCTACCGTCCCACAGATTA-3’ (forward) and 5’-TGTCGTCTCGTTTCATGCTC-3’ (reverse) were used for detecting DEC2; 5’-TTACCGGTAAGCCTAGATTAGGCC-3’ (forward) and 5’- TTGAATTCGGTAACATTATTGGTT-3’ (reverse) were used for detecting FOXC1; 5’-TCTGCACCCCCAGCAAGA-3’ (forward) and 5’-CACCCCGGAGCCTTCTGT-3’ (reverse) were used for detecting SOX4; Forward: 5’-ACGGCTTCCCAATAACAGTAG-3’ (forward) and 5’-TGTTTGACACCGAGAATTTGC-3’ (reverse) were used for detecting EZH2; Forward: 5’-CGCGATCTAAAACCACAGAAC-3’ (forward) and 5’- CAAATATGCAGCCAACACTCC-3’ (reverse) were used for detecting CDK6; Forward: 5’- AAGAAGTTCATTTACCTCCTGA-3’ (forward) and 5’- AATCACTGTCTGCTCCTTAAA-3’ (reverse) were used for detecting E2F2; Forward: 5’-CTTACAGCAGCAGGCAAAGCG-3’ (forward) and 5’- GGCTCAGGAGCTGAATGAACT-3’ (reverse) were used for detecting E2F3; 5’- TAGCCGAGGAAGAACTATGAACATAA-3’ (forward) and 5’-TGAGGTTGGTTACTGTTGGTATCATATA-3’ (reverse) were used for detecting HIF1α; 5′-CGACCACTTTGTCAAGCTCA-3′ (forward) and 5′-AGGGGTCTACATGGCAACTG-3′ (reverse) were used for detecting GAPDH; Expression of mRNA or miRNA was normalized to GAPDH or U6, respectively.

### Cell migration/invasion assay

Migration of A375 cells was assessed using Transwell chamber (Costar, Pleasanta, CA, USA) containing a polycarbonated filter with a pore size of 8 µM. A375 (3×10^4^ cells in 200 ul) cells in complete medium was mixed and the cell suspension was added to the upper chamber. The cells were allowed to migrate at 37°C incubation for 24 h and cells remaining on the topside of the transwell membrane manually removed with a cotton swab. The membrane was washed with PBS. Cells that had migrated to the underside were fixed with 95% ethanol and stained with crystal violet for counting under a light microscope. Cell migration was quantified in five random fields for each chamber.

### Luciferase reporter assay

Double-stranded oligonucleotides containing the wild-type (5’- CTAGATTTTCTTAAAAAATACCAGCAGTTACTCATGGAATATATTCTGCGTGGCCGG-3’) or mutant (5’- CTAGATTTTCTTAAAAAATGTACGTCGTTACTCATGGAATATATTCTGCGTGGCCGG-3’) sequence of the predicted miRNA-138-binding site in HIF1α 3’UTR were synthesized. Then, the wild-type and mutant 3’UTRs were cloned into a luciferase reporter vector (pmiR-REPORT, Ambion, USA) at the SpeI and HindIII site to generate Luc-HIF1α-WT and Luc-HIF1α-Mut constructs, respectively. These constructs were verified by DNA sequencing. The Luc-HIF1α-WT and Luc-HIF1α-Mut vectors were co-transfected with Lv-miRNA-138 vectors into HEK293 cells. Luciferase assays were measured using the luciferase reporter assay system (Promega, USA) 36 h after transfection. Luciferase activity was normalized by the β-galactosidase control vector activity.

### Western blot assay

The cells were lysed with RIPA buffer containing 0.1% SDS, 1% Triton-X 100, 10% glycerol, 150 mM NaCl, 0.05% Na-Doc, 5 mM EDTA (pH 8.0), 30 mM Tris-HCl (pH 7.4), protease inhibitor mixture (Thermo Scientific) and phosphatase inhibitors (Roche). After centrifugation (12000 rpm, 4°C, 15 min) to remove cell debris, the protein suspension was collected and separated by SDS-PAGE. Then, the target proteins were transferred onto PVDF membranes (Millipore Durapore, 0.45 μm pore size). The membranes were washed, blocked, and incubated with the specific primary antibodies (1:1000). The secondary antibody was IRDye 800 goat anti-rabbit or IRDye 700 goat anti-mouse (Rockland). Finally, the fluorescence signal of blots was detected using the Odyssey Infrared Imaging System (LI-COR).

### Animal study

To explore the therapeutic effect of targeting miRNA-138, female 6- to 7-week-old mice were s.c. challenged with 3 × 10^5^ LV-NC or LV-miRNA-138 overexpressing A375 cells or 3 × 10^5^ LV-NC or LV-miRNA-138 knockdown WM35 cells. Tumor growth and survival of mice were monitored every day. To explore the therapeutic effect of targeting HIF1α, female 6- to 7-week-old mice were s.c. challenged with 3 × 10^5^ melanoma cells. BAY 87-2243 were orally administered as a 2-week continuous infusion at 9 mg/kg/day (Schockel et al., 2015). Mice were monitored for tumor growth every day.

### Statistical analysis

Quantitative data were presented as mean ± SEM. All experiments were performed at least in triplicate. Differences between groups were performed by the SPSS12.0 software using two-tailed Student’s t test or one-way analysis of variance (ANOVA) for experiments with more than two groups. Survival analysis was made with the Kaplan-Meier method. Correlation analysis was performed by using two-tailed Pearson correlation coefficient. Significant differences were determined at *P* < 0.05.

## Conflict of interest

The authors state no conflict of interest.

## Author contributions

Conceptualization: H.Q., F.C.; Formal analysis: H.Q., F.C., M.C.; Writing-original draft: H.Q., M.C.; Writing-review & editing: H.Q., F.C., M.C.; Supervision: H.Q.; Project administration: H.Q.

## Funding

This work was supported by grants from the Nansha Science and Technology Project (2016MS001).

